# Early terminated transcripts and missing proteins reflect artifacts in bacterial proteomes

**DOI:** 10.64898/2026.05.19.725897

**Authors:** Giuseppe Insana, Maria J. Martin, William R. Pearson

## Abstract

The high redundancy of many bacterial proteomes can be used to evaluate proteome quality and identify sequence errors. We have used MMseqs2 clustering with subsequent filtering to identify clusters that contain sequences from at least 50% of the clustered proteomes to build sets of core proteins that include proteins from 95% of the clustered bacteria. These clusters typically capture more than 80% of proteins in the bacteria. Because these clusters have highly uniform length (the median cluster has more than 99% of its proteins at the mode length), short (<75% of mode length) or long (>133%) proteins are likely artifacts. Most “outlier” proteins are found in fewer than 10% of clusters, and “high-outlier” clusters are over-represented in a small fraction of proteomes, which often have poor proteome BUSCO fragment scores. Short-outlier proteins are artifacts; at least 80% of short-outlier genomes contain mode-length copies of the protein, which were missed because of frame-shifts, termination codons, or initiation codon choice. MMseqs2 clustering with 50% participation provides robust sets of core bacterial proteins and can be used to identify lower-quality proteomes and proteins.

## 1 Introduction

With the advent of low-cost, high-throughput, “next-generation” DNA sequencing, the number of redundant bacterial proteomes has exploded. Bacterial proteomes comprise about 51% of UniProtKB (release 2026 01) sequences, and 77% of UniParc sequences [1]. Redundant bacterial genomes have redefined our understanding of bacterial species, introducing the concepts of “open” versus “closed” pan-genomes – bacterial species that seem to accumulate more genes as additional isolates are sequenced versus species that maintain a relatively constant set of genes [2]. Indeed, the first sequencing of multiple isolates of *E. coli* [3] provided a rich catalog of examples of strain-specific genome expansion driven by horizontal gene transfer.

While genome evolution in bacterial species has been widely explored, evidence for expansion in “open” pan-genomes [4] is highly dependent on high-quality genome sequencing, assembly, and annotation that produces genuine protein sequences—sequences that are likely to be present in the living organism [2]. Spurious “bacterial” proteins have been associated with human sequence contamination [5], and the AntiFam resource has been developed to detect sequences that are often identified in organisms, but are unlikely to reflect genuine proteins [6]. Here, we describe a simple and efficient strategy for identifying a different class of spurious proteins—proteins that have been produced by fragmentation of genuine proteins, and proteins that are produced by incorrect identification of initiation or termination codons. These sequencing and annotation errors produce sequences that are either shorter, or longer, than closely related homologs. The extent to which apparent differences among bacterial proteomes represent technical artifacts remains unclear.

We have used MMseqs2 clusters to explore bacterial genome annotation quality. With ideal data, MMseqs2 would produce a set of clusters that reflects a bacterial proteome core—the proteins that are found in every species isolate. However, there are a variety of technical challenges: clustering thresholds, the cluster construction process, and the accuracy of the annotated protein predictions, that might reduce the biological reality of the clusters. In addition, there are biological challenges; some bacteria have a large range of proteome sizes, so that capturing the core proteins for a species might not be possible.

The MMseqs2 clustering strategy that we used (90% sequence identity, 50% bi-directional alignment coverage), with the requirement that a cluster contain proteins from at least half of the proteomes, produces a set of clusters that closely matches the number of genes. The requirement that each cluster contain a representative from at least 50% of the clustered proteomes actually produces clusters that typically contain representatives from more than 95% of the proteomes, with clusters containing more than 80% of the clustered proteins.

For example, an MMseqs2 clustering of 22,001 *E. coli* proteomes produces 342,137 clusters, 3,927 of which are shared by at least half of the proteomes. Indeed, more than 75% of those clusters contain proteins from more than 94% of the proteomes, Table 1 and Suppl. Table 1. The recovery of 3,927 clusters is reassuringly close to the number of proteins in *E. coli* K12 mg1655 (4,402 proteins) and the median number of proteins in our 22,001 proteomes (4,823).

**Table 1.** Bacterial proteome clusters.

| bacteria | OSCODE | omes<sup>a</sup><br>(N) | genes<sup>b</sup><br>(N) | clust.<sup>c</sup><br>(N) | cov.<sup>d</sup><br>(%) | good<sup>e</sup><br>(%) |
| --- | --- | --- | --- | --- | --- | --- |
| *E. coli* | ECOLX | 22 001 | 4 823 | 3 825 | 79.3 | 97.4 |
| *K. pneumoniae* | KLEPN | 17 924 | 5 341 | 4 552 | 85.2 | 97.1 |
| *S. aureus* | STAAU | 13 356 | 2 665 | 2 242 | 84.1 | 94.1 |
| *S. enterica* | SALER | 9 382 | 4 556 | 4 007 | 88.0 | 97.5 |
| *S. pneumoniae* | STREE | 8 865 | 2 048 | 1 701 | 83.1 | 94.6 |
| *A. baumannii* | ACIBA | 6 712 | 3 725 | 3 316 | 89.0 | 96.7 |
| *P. aeruginosa* | PSEAI | 6 367 | 6 263 | 5 406 | 86.3 | 97.1 |
| *M. tuberculosis* | MYCTX | 5 051 | 4 032 | 3 134 | 77.7 | 79.6 |
| *L. monocytogenes* | LISMN | 3 929 | 2 920 | 2 605 | 89.2 | 97.0 |
| *E. faecium* | ENTFC | 2 825 | 2 793 | 2 190 | 78.4 | 95.3 |
| *E. hormaechei* | 9ENTR | 2 540 | 4 661 | 3 812 | 81.8 | 97.2 |
| *C. difficile* | CLODI | 2 152 | 3 736 | 3 200 | 85.7 | 94.5 |
| *C. jejuni* | CAMJU | 1 845 | 1 676 | 1 440 | 85.9 | 97.0 |
| *B. pseudomallei* | BURPE | 1 632 | 5 874 | 5 015 | 85.4 | 91.5 |
| *H. pylori* | HELPX | 1 510 | 1 482 | 1 067 | 72.0 | 83.8 |
| *V. cholerae* | VIBCL | 1 484 | 3 622 | 2 676 | 73.9 | 80.5 |
| *N. meningitidis* | NEIME | 1 340 | 2 018 | 1 599 | 79.2 | 88.8 |
| *N. gonorrhoeae* | NEIGO | 1 096 | 2 078 | 1 751 | 84.2 | 90.9 |
| *L. pneumophila* | LEGNP | 937 | 3 046 | 2 460 | 80.8 | 90.1 |
| *S. flexneri* | SHIFL | 557 | 4 441 | 3 766 | 84.8 | 92.6 |
| median |  | 2 682 | 3 674 | 2 905 | 84.2 | 94.6 |

<sup>a</sup>number of proteomes clustered. <sup>b</sup>median number of genes per proteome <sup>c</sup>number of good clusters (clusters with at least 60% of proteins at the mode cluster length). <sup>d</sup>coverage of median number of genes by good clusters <sup>e</sup>yield of good clusters from the original **mmseqs2** clusters with 50% proteome coverage.

MMseqs2 cluster sets allow us to explore sources of variation in highly identical orthologous bacterial proteins. In general, proteins in MMseqs2 clusters are highly uniform in length. However, some proteomes contain dozens to hundreds of outlier proteins; proteins that are <75%, or >133% of the mode cluster length. Unusual length orthologs are surprising [7], and can contaminate efforts to characterize orthologs [8] and create biologically accurate pan-genomes and pan-proteomes. Here, we show that most of those “outlier” proteins are artifacts of genome fragmentation, sequencing errors, and mis-annotation.

Unlike eukaryotic genes, which contain introns and can be spliced into multiple isoforms [9], orthologous bacterial proteins should all have very similar lengths, particularly when they are found in the same organism. Yet some clusters constructed by MMseqs2 contained hundreds of proteins that are significantly shorter, or, to a lesser extent, significantly longer, than the mode protein length in the cluster. To examine the origin of these sequences, we clustered and analyzed redundant proteomes from 20 bacteria (Table 1) and examined their “outlier” proteins. We show that most outlier sequences are short and that in more than 80% of cases a full-length (mode-length) protein could be found in the genome of the proteome that was annotated to contain a short outlier. About 50% of the mode-length (correct) alignments in proteomes that produced short-outlier proteins contained a frameshift or termination codon. Short-outlier proteins are missing the N-terminus of the mode-length protein about 60% of the time, suggesting that initiation codon misidentification is slightly more common than incorrect termination codons. We also examined long-outliers and found that more than 80% of long-outliers were produced by annotated open reading frames (ORFs) that spanned two adjacent genes.

Outlier frequency is highly correlated with Proteome BUSCO fragment score [10, 11], and less well correlated with other BUSCO quality measures. However, we found several reference proteomes with very low BUSCO fragment scores but relatively high cluster outlier content. Thus, outlier content provides an additional measure of proteome quality.

## 2 MATERIALS AND METHODS

### 2.1 Proteome clustering with MMseqs2

The 20 bacterial species with the largest number of non-excluded proteomes in the UniProt and UniParc protein sequence repositories were downloaded as individual proteome files in fasta format from UniParc. We excluded proteomes associated with genome assemblies that have been flagged by either UniProt or RefSeq quality control analyses with warnings [12], and proteomes that contained fewer than 200 proteins. The names of the bacteria and associated OSCODEs are shown in 1. and the NCBI TaxonID’s, number of proteomes, and the number of sequences are shown in Suppl. Table 1. The full list of all the proteomes analyzed, together with their assembly accessions, can be found in the Supplementary Data (see Data Availability).

The protein sequences from each bacterial species were clustered using MMseqs2 (`mmseqs easy-cluster` with parameters cov-mode 0, seq-id-mode 0, min-seq-id 0.9, coverage 0.5, cluster-mode 0, clust-hash 1, alignment-mode 3, kmer-per-seq 100, max-seqs 10000, s 7.5 and substitution matrix VTML10), labelling the protein identifiers in each cluster with their corresponding proteome identifiers. 90% sequence identity was chosen for clustering proteomes from the same bacterial species; the actual median sequence identity in the cluster is much higher [13]. The number of proteomes, the initial number of proteins, the percentage of clusters with representatives from > 1 proteome, ≥ 10% of clustered proteomes, or ≥ 50% percent of clustered proteomes clusters, are shown in Suppl. Table 1. Only clusters containing proteins from at least 50% of the clustered proteomes were examined. Reducing the proteome participation level from 50% to 10% increases the number of clusters about 1.5-fold, but increases the number of proteins only about 10%. A `nextflow` pipeline with `python` scripts was used to cluster and filter the proteomes. This is available at https://github.com/g-insana/ProteomeCluster.

European Nucleotide Archive (ENA, [14]) Swagger APIs were used to retrieve bacterial genomic sequences and related metadata (endpoints `fasta/`, `summary/`, `xml/` and `embl/` of https://www.ebi.ac.uk/ena/browser/api/).

### 2.2 Identification of protein length outliers

Lists of MMseqs2 clusters were parsed (1) to extract the proteins and their lengths from each cluster, (2) to identify the mode protein length, and (3) to extract the members of the clusters that were either <75% or >133% of the mode protein length. The number of outlier sequences, and the average and median number of outliers per cluster are shown in Suppl. Table 2. Since the number of proteomes ranged from 557 (*S. flexneri*) to 22,001 (*E. coli*, Table 1 and Suppl. Table 1), we identified two sets of proteomes: (1) proteomes with the most clusters with outlier proteins and (2) a random sample of up to 2,000 proteomes with outliers. We also sampled up to 100 short-outlier proteins and their corresponding proteomes, for use in translated alignments, as well as a set of long-outlier proteins and their corresponding mode-length protein genomes, to determine whether a long protein could be found in a genome that produced a mode-length sequence.

A small subset of clusters (typically < 5%, Table 1), produced many outliers (sometimes 50% or more of the proteins in the cluster) because the cluster protein length distribution did not have a characteristic mode length. We excluded these clusters from our analyses, and focused on clusters where at least 60% of the members of the cluster were exactly the mode protein length (Table 1).

### 2.3 Protein-DNA alignment with tfastx

To test whether genomes annotated to contain short-outlier proteins might actually contain full-length (mode-length) proteins, short-outlier genomes were searched with mode-length (full-length) proteins using the `tfastx` program from the `FASTA` package (version 3.8, released July 2025). The `tfastx` program aligns a protein query sequence to a DNA sequence database (in this case, a bacterial genome), allowing alignments to span across termination codons and frameshifts [15, 16], using a scoring matrix (`-s MD10`) designed to identify sequences sharing ≥ 90% identity [17]. The analysis script identifies up to 100 proteomes, each of which produces up to 20 short-outlier proteins and their associated clusters. Mode-length (full-length) protein sequences were then aligned with short-outlier genomes with `tfastx`. For bacteria with smaller numbers of proteomes, or with smaller numbers of short-outliers, fewer searches were done. On average, ∼2000 `tfastx` alignments were done to search for full-length alignments in short-outlier genomes (∼1500 for the worst clusters, and ∼500 for the sampled clusters).

Alignments were reported using blast-tabular format (`-M8CBl`), and the alignment summaries were parsed to identify alignments that spanned at least 95% of the length of the query sequence (the longer mode-length sequence for short-outliers, or the long outlier sequence for long-outliers). A single alignment with at least 90% identity was counted as full length if it extended to at least 95% of the query sequence length (the long sequence), or if a set of ≥ 90% identical alignments that overlapped by less than 5% of the query sequence length added up to at least 95% of the query sequence length (a multiple-hit alignment).

To determine whether a longer protein extended in the N-terminal or C-terminal direction, outlier proteins were aligned to mode-length proteins using the `ssearch` program from the `FASTA` package using the MD10 scoring matrix.

### 2.4 Statistical analysis and plotting

Graphical figures were constructed using the `ggplot2` library and `R` version 4.5.1 (2025-06-13) [18, 19].

The scripts used to cluster and parse the cluster data, perform `tfastx` and `ssearch` alignments, parse alignment results, and prepare final figures are available from GitHub: github.com/g-insana/ett_ms.

## 3 RESULTS

With the dramatic decrease in sequencing costs over the past 15 years, the number of sequenced bacterial genomes has exploded, producing more than 400,000 *E. coli* genomes, and tens of thousands to thousands of genomes for many other widely studied bacteria [20]. Although not all sequenced genomes contain protein annotations, the 2026 01 release of UniProtKB (28-January-2026) contains 9 reference *E. coli* proteomes, and another 255 other *E. coli* proteomes whose proteins are included in UniProtKB. In addition, the more comprehensive UniParc database contained 18,735 “redundant” *E. coli* proteomes, and 124,444 “excluded” proteomes.

These large numbers of “duplicate” proteomes from the same bacterial species allow us to explore the uniformity of virtually identical proteins, and, when orthologous protein sets are highly uniform, to use non-uniformity to identify proteins, and the proteomes that contain them, that are likely to reflect sequencing, assembly, and annotation errors. To characterize bacterial protein set uniformity, we have used MMseqs2 to cluster proteomes from the same species under conditions that produce numbers of clusters that closely match the number of proteins in each species, after filtering the clusters to require that each cluster contain proteins from at least 50% of the clustered proteomes.

### 3.1 MMseqs2 clustering identifies bacterial protein cores

MMseqs2 clusters can be examined from two perspectives: (1) how well do the clusters capture the diversity of proteins in the proteomes that were clustered, and (2) how many of the proteomes are included in each cluster. In a hypothetical ideal case, where every proteome had exactly the same set of proteins, there would (1) be one cluster for every single-copy gene in proteome, and (2) each cluster would contain a gene from each proteome included in the clustering. So, for example, if 10,000 proteomes from a bacteria with 4,000 genes were clustered, there would be 4,000 clusters, each of which contained 10,000 proteins.

n

We used MMseqs2 in a relatively conservative mode (–cov-mode 0), requiring 50% coverage of the longer of the query and the subject sequences, with 90% identity in the cluster (–min-seq-id 0.90). The resulting MMseqs2 clusters were then filtered to select clusters that contained proteins from at least 50% of the clustered proteomes. Although the filter only required clusters to contain 50% of the proteomes, half of the clusters contained proteins from more than 99.1% of the proteomes (Fig. 1), and the 5.6% of the complete MM-seqs2 cluster set that contain proteins from 50% of the proteomes include 85% of the proteins that were clustered (Suppl. Table 1, over-all median). Across the 20 bacterial species we examined, MMseqs2 clusters capture a very high fraction (from 72%, *H. pylori* to 89.2%, *L. monocytogenes*) of the median number of proteins found in each of the bacterial species (Table 1, “cov.” column, see also Suppl. Table 1 sequences ≥ 50%). Thus, MMseqs2 / 50% proteome participation clusters typically capture more than 85% of protein diversity of the clustered proteomes—capturing most of the protein core. In addition, more than 99% of the clustered proteomes contribute to those core protein clusters.

**Figure 1.**
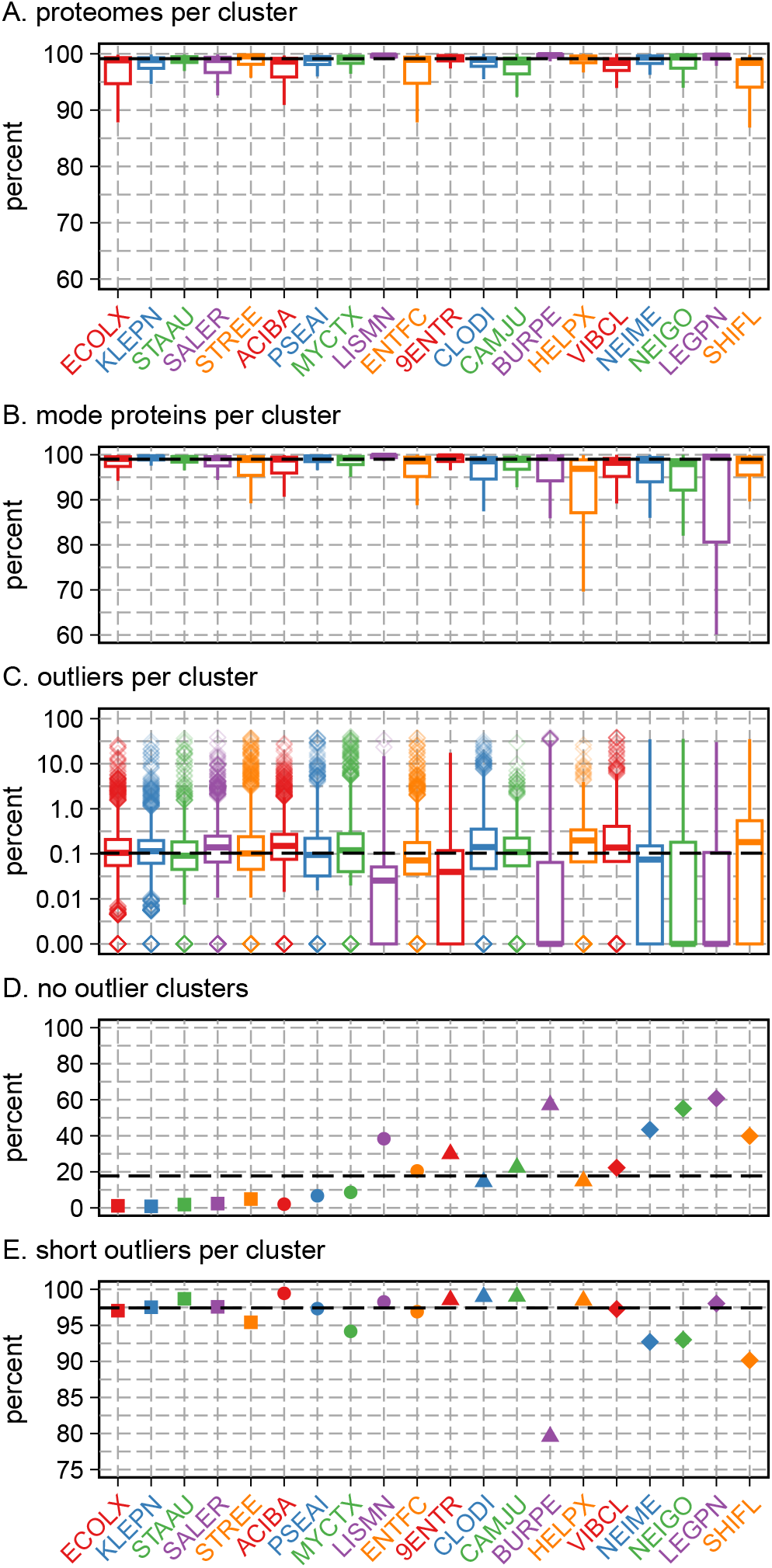
Cluster uniformity, quality, and error types, across the 20 bacterial species. (A) Percent of all proteomes (including proteomes not represented in the cluster) included in a cluster. About 10% of clusters are excluded from this plot because the fraction of proteins/cluster was between 50% and 60%. (B) percent of proteins in cluster at mode length. (C) The distribution of outliers from each cluster in a bacterial species. Cluster numbers range from 1,067 (HELPX, *H. pylori*) to 5,406 (PSEAI, *P. aeruginosa*), with a median of 2,905 and first and third quartiles of 2,080 and 3,673. The number of clusters for each bacterial species is shown in Suppl. Table 1, and the number of outliers in Suppl. Table 2. (D) Percent of clusters in each organism that had no outliers. (E) Percent of outliers that are shorter than 75% of the mode length. Dashed lines indicate the medians of the medians across the 20 bacteria. **Alt text:** Five panels labelled A to E showing boxplot distributions (A to C) or plotted values (D and E) for the 20 bacterial species examined. In each panel a dashed line indicates the value of the median of medians.

While the number of MMseqs2 clusters created with the 50% proteome coverage filter closely matches the median number of proteins in the set of proteomes we clustered (Suppl. Fig. 1A), and most clusters contain proteins from more than 97% of the proteomes that were clustered, these matching numbers do not guarantee that the proteins in the cluster are the same as the proteins widely shared across proteomes from the same bacterial species (pan-genome proteins). To confirm that the MMseqs2 50% proteome coverage clusters correspond to the shared pan-genome, we compared a set of *E. coli* pan-genome proteins identified by Horesh *et al*. [21] to a set of proteins constructed by selecting a single mode-length protein from each of the clusters from the `ECOLX` proteome set (the 3,927 clusters include 102 clusters that did not meet the >60% of proteins at mode length shown in Table 1). When the 3,825 `ECOLX` clustered proteins were compared to the 3,932 pan-genome proteins identified by Horesh *et al*. that are shared by at least 50% of lineages, 95.8% of the mode `ECOLX` proteins share a ≥90% identical match over ≥90% of the alignment, while 96.6% of the pan-genome proteins share ≥90% identity over ≥90% of the alignment. Thus, the MMseqs2 50% proteome coverage protein set is ∼95% identical to a core pan-genome *E. coli* protein set, confirming that the MMseqs2 clusters produce a comprehensive and very complete sample of shared *E. coli* pan-genome proteins.

### 3.2 Identification of “missing” proteins with tfastx

While the 50% proteome participation clusters include proteins from 99.1% of the proteomes at the median, it is unclear whether the proteomes that did not contribute to clusters actually lacked those proteins, or if they were “missing” because of assembly and annotation artifacts. To search for the “missing” proteins, we (1) identified proteomes that contributed to fewer than 2/3 of clusters for the 20 bacterial species, (2) randomly selected 20 proteomes from the low contributing proteomes, (3) randomly selected up to 50 mode-length proteins with lengths >100 amino acids from the clusters that were missing from the proteomes, and (4) used `tfastx` to align the mode-length proteins to the genomes that were annotated to be missing a member of the cluster.

Across 19 bacterial species, `tfastx` searches with a “missing” mode protein produced a full-length alignment 95.2% of the time (Suppl. Fig. 2A, one bacteria, *N. gonorrhoeae* did not have any proteomes that contributed to fewer than 2/3 of the clusters). Moreover, unlike the recovery of mode-length proteins in proteomes annotated to produce short proteins below, almost all the proteins were missed because of frameshift errors (median 96.8%, Suppl. Fig. 2B). These proteins could have been missed because they were not included in the 50% participation clusters, but searches with the missing mode-length proteins identified a homolog in the corresponding proteome less than 2% of time, and those few homologs were more than 90% identical only about 18% of the time. Thus, proteomes that appear to be missing proteins are likely to contain them; they were missed because of sequencing or assembly errors. *S. pneumoniae* (STREE), *M. tuberculosis* (MYCTX), and *S. flexneri* (SHIFL) are exceptions; for these bacteria, the “missing proteins” were only recovered between 26% (STREE, SHIFL) and 38% (MYCTX) of the time; >90% identical orthologs were recovered >80% of the time (median 95.2% across 19 bacteria) from the other 16 bacteria with missing cluster members.

**Figure 2.**
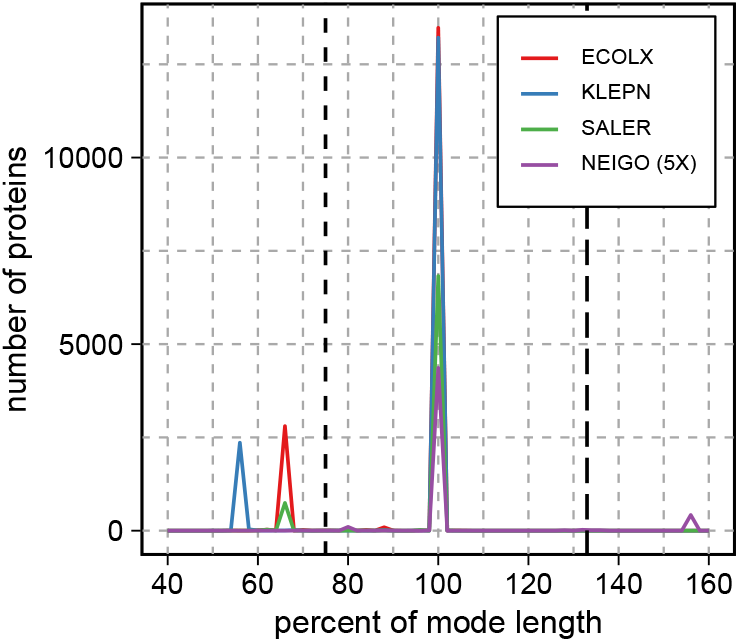
Distribution of protein lengths, plotted as percent of the mode protein length, for one cluster from each of four different bacteria: *E. coli* (ECOLX), *K. pneumoniae* (KLEPN), *S. enterica* (SALER), and *N. gonorrhoeae* (NEIGO). NEIGO protein numbers were multiplied by 5 for visibility. Vertical lines indicate the limits (*<*75% and *>*133%) used to classify outliers. 17.4% (ECOLX), 15.6% KLEPN, and 10.3% (SALER) of the proteins in each cluster were labeled as outliers, placing these clusters among the worst 0.2% of all clusters for each bacteria; 8.0% of the NEIGO proteins in this cluster are long-outliers, among the worst 3% of clusters (overall, 75% of NEIGO clusters contain short outliers). **Alt text:** Line graph showing the distribution of protein lengths as a percentage of mode length for single protein clusters across four bacterial species.

### 3.3 MMseqs2 clusters have uniform lengths

MMseqs2/50% participation clusters can be used to identify incorrect protein sequences because orthologous (typically >95% identical) proteins from the same bacterial species have highly uniform lengths. Although our MMseqs2 parameter settings required an alignment to include only 50% of the longer protein to be included in a cluster (thus, both a 100 residue protein and a 400 amino-acid protein that aligned to a 200 cluster representative with 90% identity could be included in the same cluster), almost all (97.1% across the 20 bacteria examined) proteins in the clusters had exactly the same length (the mode length, Fig. 1B). Clusters that had fewer than 60% of the proteins at the mode length occurred less than 5.5% of the time (median over all 20 bacteria, Table 1). More inclusively, 75% of the proteins in a cluster (the bottom of the boxes in Fig. 1B) had the mode length 95% of the time.

Thus, MMseqs2 –cov-mode 0 -c 0.5 (50% coverage) –min-seq-id 0.9 (90% identity), with the requirement that clusters contain proteins from 50% of the proteomes, produces a set of clusters that closely matches the median number of proteins in the clustered proteomes, and 95% of the proteins in those clusters are exactly the same length more than 95% of the time. These MMseqs2 parameters, with filtering for 50% proteome inclusion, produce comprehensive and uniform protein sets.

### 3.4 Some clusters have many “outlier” proteins

While virtually all the proteins in most clusters have the same length (Fig. 1B), some clusters contain large numbers of “outliers”. We have defined “outliers” as proteins with lengths either <75% or >133% of the cluster mode length. We chose the <75%/>133% thresholds to count only dramatic length variation; many clusters have non-mode-length proteins that are do not meet the <75/>133 threshold (Suppl. Fig. 3). Fig. 2 shows four distributions of normalized protein lengths from a single cluster from each of four bacteria, *E. coli* (ECOLX), *K. pneumoniae* (KLEPN), *S. enterica* (SALER), and *N. gonorrhoeae* (NEIGO). These clusters were chosen because they have a high frequency of outliers (8–17%); in most clusters, fewer than 0.1% of the members are outliers (Fig. 1C).

**Figure 3.**
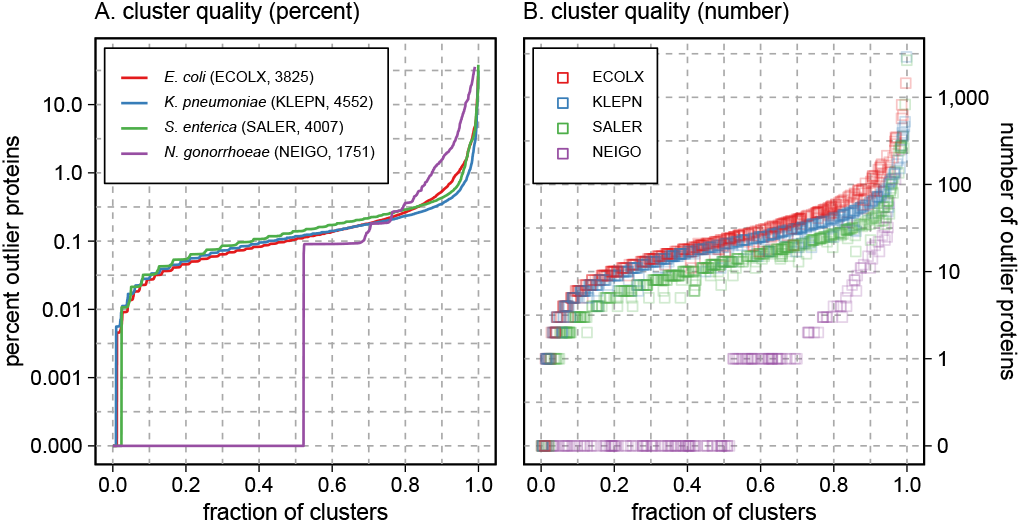
Distribution of outliers in protein clusters for four bacteria: *E. coli, K. pneumoniae, S. enterica* and *N. gonorrhoeae*. Protein clusters are sorted from fewest to most outliers (A) distribution of cluster quality showing the percent of outlier proteins in the cluster. (B) distribution of outlier proteins by number. **Alt text:** Dual panel graph showing the percentage and absolute number of outlier proteins across sorted protein clusters for four bacterial species.

The overall length distributions of all the clusters in the four bacteria shown in Fig. 2 are included in Fig. 1B, which also summarizes how often a cluster has outliers (Fig. 1C). The absolute numbers of outliers for each of the 20 bacterial species is shown in Suppl. Table 2, which also displays the average and median number of outliers per cluster. Fig. 1 also displays how often clusters have no outliers (panel D), and how often outliers are short (<75% mode length, panel E) for all twenty bacteria. Outlier proteins are relatively rare (Fig. 1, Suppl. Fig. 1); the median percentage of outliers per cluster is around 0.1% (Fig. 1C), or about 20 proteins for *E. coli* (ECOLX), where 22,001 proteomes were clustered, and only 1 or 2 proteins in the bacteria with fewer than 2,000 clustered proteomes. Three bacteria, *B. pseudomallei* (BURPE, 1,632 proteomes), *N. gonorrhoeae* (NEIGO, 1,096 proteomes), and *L. pneumophila* (LEGPN, 937 proteomes) have no outliers in more than half of their clusters (Fig. 1D). Most outlier proteins (97% of outliers, Fig. 1E) are shorter than the mode length in a cluster. *B. pseudomallei* (BURPE) has the largest fraction of long outliers—about 23% of clusters with outliers have long outliers—but only 40% of clusters in this bacteria had any outliers (Fig. 1C).

The outlier frequencies summarized in Fig. 1C and D reflect the number of short (<75% mode-length) and long (>133% mode-length) outliers. We chose these thresholds to be conservative; proteins that are slightly longer or slightly shorter than the mode length are not considered outliers. Changing the thresholds to be less stringent, e.g. outliers are <90% or >111% of mode length doubles the number of outliers (the median fraction of outlier clusters across the 20 bacterial species increases from 0.104% to 0.223%), but does not change the outlier pattern across the 20 species. Likewise, making the outlier thresholds more stringent (<66%, >150%) decreases the median fraction of outliers to 0.051%, again, without changing species patterns.

### 3.5 Most outlier proteins are concentrated in a small fraction of clusters

Fig. 3 plots the percentage of outlier proteins per cluster for every cluster for the four bacteria shown in Fig. 2. Ninety percent of the proteomes have fewer than 0.2% (KLEPN)– 0.4% (ECOLX, SALER) of clusters with an outlier protein. Each proteome contributes a single protein to each cluster, so, for *E. coli*, with 3,825 clusters, half the proteomes (∼11,000) have ≤3 outlier proteins, 90% have fewer than 7, while the worst proteomes have more than 200.

Although the frequency of outliers in most clusters follows linear exponential, a small fraction of clusters in each of the bacteria account for half of the total number of outliers. In Fig. 3, which excludes clusters with <60% of mode length proteins, 50% of the total outliers are found in the worst 4.0% (ECOLX), 6.7% (KLEPN), 4.6% (SALER), and 1.7% (NEIGO) of clusters. Thus, most outlier proteins are produced by a relatively small fraction of clusters.

### 3.6 Most outlier proteins are concentrated in a small fraction of proteomes

Just as we can examine the distributions of protein numbers, protein lengths, and outlier frequencies across protein clusters (Fig. 1), we can also ask how these statistics are distributed across proteomes. Suppl. Fig. 1 shows how our MM-seqs2 clustering with 50% proteome participation captures the proteins in the 20 bacteria. While Fig. 1 shows the distribution of protein numbers, lengths, and outliers across the ∼3,000 clusters produced from each bacteria, Suppl. Fig. 1 summarizes these values across the 22,001–597 proteomes from each bacteria. For example, in Fig. 1A, the number of proteomes per cluster is divided by the total number of proteomes for each cluster. In Suppl. Fig. 1A, the total number of clusters that contained each proteome is divided by the total number of clusters produced from each bacteria. Thus, for *E. coli*, half of the proteomes are included in 91% of the clusters, and 75% of the proteomes contribute to 88% of the clusters. Likewise, most of the proteomes (98%) have the mode protein length for the clusters that they belong to, confirming that most proteins in both clusters and proteomes have uniform lengths. Across the distribution of proteomes, there is a somewhat higher median outlier frequency (0.2%) than across clusters, but in both cases, the frequencies are quite low at the median. As seen by the extreme values (diamonds in Suppl. Fig. 1C), some proteomes have outliers from 10% to more than 30% of the clusters they contribute to.

The relationship between cluster quality and proteome quality is shown in more detail in Suppl. Fig. 3. Here, the top three panels show the sample proteome quality shown in Fig. 3, while the 10 numbered panels below them show the distribution of protein lengths for 10 high-outlier clusters from each of the bacteria. If high-outlier clusters were always produced by high-outlier proteomes, then the red + symbols, which indicate outliers, would largely be found to the right of the proteome axis, because they were found in the proteomes that had the most outliers. This is seen in the A panels (A.4, A.7, A.10), less frequently in the B panels (B.8, B.9), and more frequently in the C panels (C.2, C.6-C.9). In addition, Suppl. Fig. 3 shows that not all non-mode-length proteins have been classified as outliers: 2–4 clusters from each bacteria have many + symbols—proteins that are not mode-length, but not different enough to be classified as outliers. Suppl. Fig. 3 also shows that non-mode-length proteins have strongly preferred lengths. Outliers and other non-mode lengths have characteristic lengths.

### 3.7 Outliers and proteome quality

If, as the uniformity of mode-lengths suggests, outliers are not the result of genuine biological protein length variation, and if they are not uniformly distributed across proteomes, then outlier distributions may reflect some property of the genome assembly. Fig. 4B, D, and F plot the relationship between outlier frequency and a widely used measure of proteome quality, the Proteome BUSCO fragment score [10]. Because of the wide dynamic range of outlier frequency and BUSCO fragment scores across the three bacterial proteome sets in Fig. 3, the relationship is plotted on a log-log scale, but the correlation coefficients are calculated for untransformed outlier frequencies and BUSCO fragment scores. Outlier frequency is strongly correlated with BUSCO fragment score, though for all three bacteria, there is a large range of outlier frequency associated with a BUSCO fragment score of 0. The BUSCO score focuses on a set of core enzymes expected to be found in all organisms, while the outlier frequency looks at 80+% of the proteins in each bacteria. The outlier frequency appears to capture quality issues that are not found by BUSCO.

**Figure 4.**
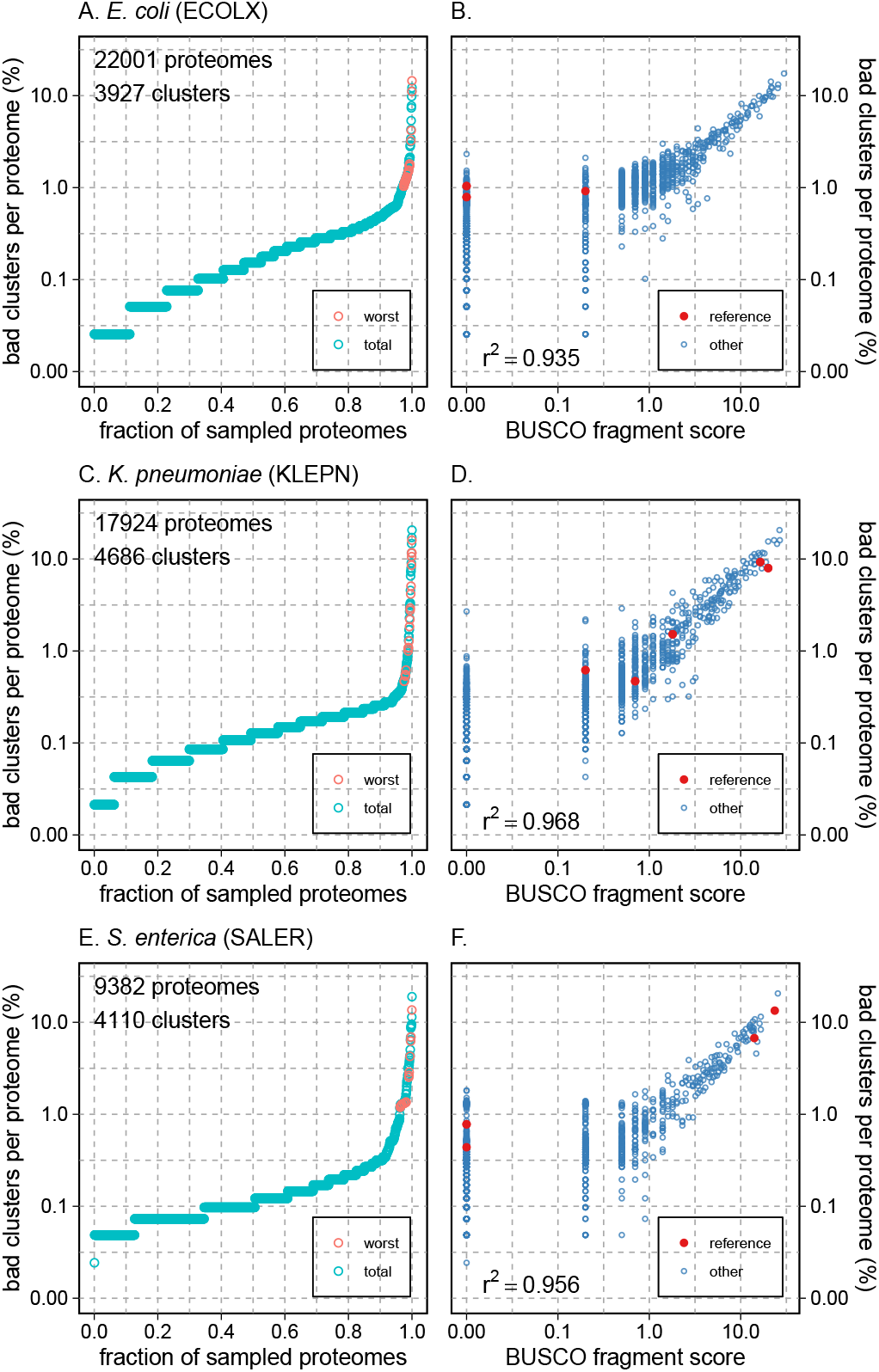
Distributions of outliers in three proteomes. Panels (A), (C), and (E) show the percent of clusters with outliers in the proteome, ranked from the fewest outliers to most. (A) ECOLX, the organism with the largest number of proteomes, has proteomes that range from fewer than 0.03% bad clusters (one bad cluster per proteome) to more than 10% of clusters. Pink circles show a sample of proteomes with the largest number of clusters with outliers; blue circles are randomly sampled from the entire proteome set. (B, D, F) Correlation of bad cluster frequency with Proteome BUSCO fragment score. Reference proteomes (those distributed in the UniProt Reference Proteome set) are shown as solid red circles; other proteomes are shown with open circles. Additional BUSCO quality measures are plotted in Suppl. Fig. 4. **Alt text:** Six panels displaying the percentage of bad clusters per proteome and their correlation with BUSCO fragment scores for three bacterial species.

The different colored circles in Figs. 4B, D, and F denote two classifications of proteomes in UniProtKB and UniParc. The red colored “reference” proteomes are included in both UniProtKB and the UniProt Reference Proteome set [22]. The “other” proteomes are included in UniProtKB (these non-reference proteomes will be dropped from UniProtKB in release 2026 02, but will be available from UniParc).

BUSCO analysis provides other measures of proteome quality, including BUSCO completeness (combined and single), and BUSCO missing scores [10]. These alternate measures of proteome quality also correlate with outlier frequency, but much less dramatically (Suppl. Fig. 4).

### 3.8 Full-length proteins can be found in proteomes annotated to produce short outliers

To explore whether short-outlier proteins are technical artifacts of genome assembly and annotation, we took mode-length (full-length) proteins from clusters with either large percentages of short outliers (Fig. 6, red symbols) or randomly sampled across the set of clusters for a bacteria (cyan symbols) and compared those proteins to the genomes that produced the short-outlier proteins, using the `tfastx` alignment program. `tfastx` aligns a protein sequence to a DNA sequence, allowing alignments to extend across frameshifts and termination codons [15, 16]. When it was first described, `tfastx` was shown to be able to extend protein sequence alignments in genome annotations of 24 genes in *H. influenzae* based on very statistically significant similarity (better than E()< 10^−10^) to *E. coli* or other *H. influenzae* proteins, so we were not surprised when `tfastx` produced mode-length alignments in genomes annotated to contain short outlier proteins.

`tfastx` can produce full-length alignments of mode-length proteins with short-outlier genomes two ways: either (1) as a single alignment that aligns the full-length query protein from beginning to end (Fig. 5A,B), or (2) as several alignments, which cover the full length of the query protein (Fig. 5C,D). Suppl. Fig. 5 shows residue alignments for three additional comparisons of mode-length proteins to short-outlier genomes.

**Figure 5.**
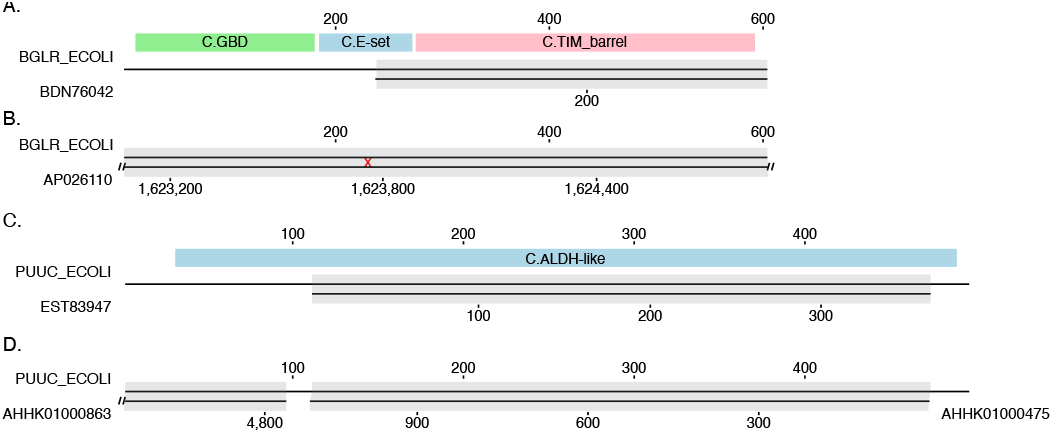
`tfastx` alignment of full-length proteins to genomes that produced short-outliers. The `ssearch` and `tfastx` programs were used to align full-length (mode-length) proteins to either (A,C) short-outlier proteins from the same cluster, or (B,D) the bacterial genome sequences that produced the short outliers. Solid black lines represent the subject and query sequences. The aligned regions are highlighted in grey. (A) Alignment of `BGLR ECOLI`, *E. coli β*-glucuronidase, to the short-outlier `BDN76042`, which was produced by the *E. coli* genome `GCA 025996495.1` with the `ssearch` program. Also shown are the `Pfam` domains annotated on `BGLR ECOLI` . (B) Alignment of `BGLR ECOLI` to `GCA 025996495.1/AP028110`, the genome that produced `BDN76042`, using `tfastx` . `GCA 025996495.1` contains a single 5,395,638 nt sequence entry (`AP026110`). The red x indicates a frameshift. The 603 amino acid full-length alignment of `BGLR ECOLI` to `AP026110` is 99.3% identical (4 conservative substitutions). (C) Alignment of `PUUC ECOLI`, *E. coli* NADP/NAD-dependent aldehyde dehydrogenase, to a short-outlier from the cluster, `EST83947` . The `Pfam` domain annotation for `PUUC ECOLI` is shown in blue. (D) Alignment of `PUUC ECOLI` to the genomic DNA that produced `EST83947`, `GCA 000498175.1/AHHK01000475` . The `GCA 000498175.1` genome contains 4654138 nucleotides in 1173 sequences, and `PUUC ECOLI` is 100% identical across translations of two contigs from that genome. **Alt text:** Structural diagrams showing protein-to-genome alignments comparing full-length sequences to short-outlier annotations.

Single contig full-length alignments happen most frequently when `tfastx` aligns against long genomic regions (Fig. 6, circles), while multiple overlapping alignments are seen when aligning to shorter genomic fragments (Fig. 6 “+” symbols).

While different bacterial proteome sets show a range of mode-length protein recovery from short outliers, mode-length proteins are found in the proteomes that produced short outliers about 80% of the time on average (Suppl. Fig. 7A). About 60% of the short outlier genomes contain a one-contig alignment that produces a full-length protein (Suppl. Fig. 7B). Almost half of short-outliers are produced by frameshifts or termination codons produced by sequencing errors (Suppl. Fig. 7C,D). The remaining short-outlier alignment errors do not have sequencing errors; for one-contig alignments they must reflect the incorrect choice of an initiation codon (Suppl. Fig. 7C). This observation is supported by the frequency of N-terminal extensions in the genomes that produced short proteins (Suppl. Fig. 7E). Short outlier proteins, the most common types of outliers by far, are artifacts produced by sequencing errors, gaps in genome assemblies, and annotation errors that select an incorrect initiation codon.

**Figure 6.**
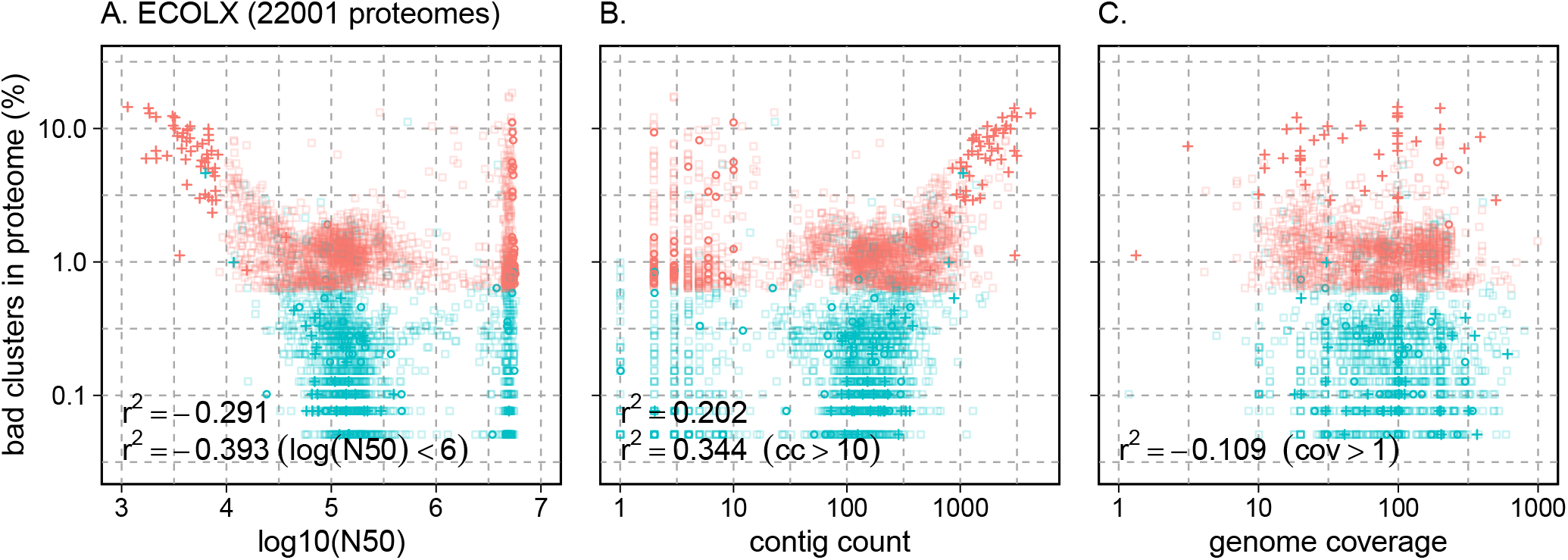
Genome assembly quality and high-outlier proteomes. A sample of the 22,001 *E. coli* proteomes with the highest fraction of outlier proteins (red) or a random sample of proteomes (blue) are plotted against three measures of genome assembly quality. Up to 2,000 of the mode-length proteins were aligned with to the outlier proteomes using `tfastx` ; alignments that produced full-length (mode-length) proteins from a single contig (circles) or multiple contigs (‘+’) are shown. A larger set of proteins that was not aligned is plotted with squares. (A) Comparison of cluster quality and log(N50), where N50 is the median length of contigs for the assembly. (B) Cluster quality versus the number of contigs in the assembly. (C) Cluster quality versus genome coverage. Additional proteomes are plotted in Suppl. Fig. 6. **Alt text:** Three scatter plots comparing *E. coli* genome assembly quality measures with the percentage of bad clusters in the proteome.

### 3.9 Long outlier proteins can also be found in proteomes that produce mode-length proteins

Just as genomes that produce short outlier proteins contain gene regions that can produce a full-length protein, genomes that produce mode-length proteins contain gene regions that could potentially produce a long-outlier protein. Long-outlier proteins are much rarer than short-outliers (short outliers account for more than 95% of length outliers, Fig. 1E). But when long outliers occur, we can ask whether a genome that produced a mode-length protein might also encode a long-outlier, by searching the mode-length genome with the long-outlier protein using `tfastx`. Just as more than 80% of short-outlier genomes contain a genomic region that could encode a mode-length protein (Suppl. Fig. 7A), most mode-length genomes have the potential to produce a long-outlier protein. In general, long-outlier alignments are typically of lower quality than the mode-length alignments seen with short outliers; they tend to have more gaps, and many of the long proteins have runs of `XXXX`.

To understand better whether long outliers are likely to be genuine proteins, we searched a comprehensive bacterial protein database with 20 long-outlier extensions from *E. coli* and 20 from *B. pseudomallei* (the bacteria with the largest fraction of long outliers) and compared the number of close homologs ((*E*() < 10^−10^, identity > 50%) found with the extended sequence to the number found after a search with the comparable length region of the mode-length sequence. While searches with the protein fragment from the mode-length protein found high identity, full-length alignments in every case, searches with the portions of the long-outlier proteins that were not found in the corresponding mode-length protein produced two classes of results. About 10% of the time, the extended protein sequence did not produce a statistically significant (*E*() < 10^−10^ result with any bacterial proteins. The remainder of the time, the extended-fragment searches produced high identity, high significance alignments that did not include either the first 20–50 amino-acids at the N-terminus of the proteins (for extended regions from at the C-terminus of the mode-length protein), or for the last 20–50 amino-acids from the C-terminus of extended sequence (for extended regions from the N-terminus of the mode-length protein). These alignments were consistent with the long-outlier protein including a portion of the protein either up-stream or down-stream from the mode-length protein. Long-outlier proteins do not exist *in vivo*; they simply reflect the ease with which open reading frames can be extended through intergenic regions in high-coding density bacterial genomes.

## 4 DISCUSSION

We have used MMseqs2 clustering to efficiently identify bacterial pan-genome cores, which in turn can be used to quantify the overall quality of a bacterial protein set, and to identify individual protein sequence artifacts. Orthologous, highly identical, proteins sequenced from the same bacterial species can be readily identified with MMseqs2 clustering when 50% proteome participation is required. These clusters appear to approximate protein cores for the species, both in the number of clusters produced and the number of distinct proteomes they include. The lengths of the proteins in these clusters are highly uniform, with 97% (median, 95% first quartile) of the proteins in a cluster having the mode length (after excluding the small fraction— ∼5% of clusters where 60% of the members do not share the mode length). Because they are rare, deviations from the mode length in a cluster can be used to characterize proteome quality and protein accuracy.

Because the MMseqs2 50% participation clusters are both comprehensive and uniform in length, we were surprised to find clusters with large numbers of proteins with unusually short (<75% of mode) or long (>133% mode) lengths—*i*.*e*. “outlier proteins”. Outlier proteins are rare; 50% of clusters contain fewer than 0.1% outliers (Fig. 1C, 22 proteins per cluster for *E. coli*, with 22,001 proteomes; less than 1 protein per cluster for *S. flexneri*, with 557 proteomes), and three bacteria have no outliers in more than half their clusters (*B. pseudomallei, N. gonorrhoeae*, and *L. pneumophila*). The very high length uniformity in these clusters suggests that outliers are artifacts.

Most outliers are short. In 50% of clusters, 97% of outliers are short across the 20 bacteria, and even in *B. pseudomallei*, which had the smallest fraction of short outliers, 79.6% of the outliers were short. And short-outliers are artifacts. In 80% of the short-protein producing genomes that we examined, a `tfastx` alignment of the mode-length protein produced a full-length alignment. The 80% recovery rate for full-length proteins is conservative; additional full-length proteins are likely present but were not found, either because the contigs that encode them were too short, or were not sequenced. Cryptic full-length proteins explain why most outliers are short, and why high frequencies of outliers are associated with lower quality proteomes/genomes.

We cannot know for certain that short-outliers are sequencing, assembly, or annotation artifacts; it is possible that the frame-shifts and termination codons that produced the short proteins reflect genuine evolutionary divergence or pseudogenes. We believe the biological explanation is less likely because of the high identity of the alignments between the full-length (mode-length) proteins and the short-outlier genomes. The average and median `tfastx` alignment identities across the 18 bacterial species with short outliers was 98.2%. *H. pylori* shared the worst average `tfastx` alignment identity, 96.6%, a value that is much higher than the 90% identity required for a full-length alignment to be scored. This high identity is more easily explained by technical artifacts, which do not reflect evolutionary distance. Short outliers are a genome assembly/annotation artifact that can both decrease the apparent size of a bacterial pan-genome core, and increase the number of proteins outside the core.

We believe that our central results, that MMseqs2 clustering with 50% participation produces a biologically accurate set of core proteins, and that sequence length outliers can be used to identify and correct protein artifacts, are robust to the clustering and threshold parameters used in this study. We are comfortable that modest changes in the MM-seqs2 clustering would have small effects, because we produce clusters that have much higher proteome participation (∼99%) than required (50%). The absolute numbers of outliers do depend on the outlier thresholds. As expected, less stringent thresholds produce more outliers, and more stringent thresholds produce fewer outliers, but the overall patterns of outliers across the bacterial species remains consistent. Bacterial species with proteomes with many outliers (`KELPN`, `SALER`) have higher numbers of outliers at more and less stringent thresholds, while proteomes with fewer outliers (`BURPE`, `LEGPN`) have fewer clusters at the other thresholds. Indeed, we can imagine that more stringent outlier identification would improve proteome accuracy.

High-outlier frequencies reflect lower proteome quality, which in turn reflect lower genome assembly and annotation quality. Identification of protein sequence length outliers complements the widely used BUSCO measures of proteome quality[10], focusing on highly conserved orthologous protein length, rather than the presence (or absence) of core pathway proteins. Mode-length outlier identification exploits the very high identities of orthologs from the same species, and thus makes protein specific error predictions as well as providing an overall measure of proteome quality. However, accurate orthologous protein length measurements require many dozens of proteomes from the same species, unlike BUSCO, which can evaluate the protein set from a single proteome. Currently, in UniProt, there are almost 200 bacterial species with more than 100 proteomes, which can be scanned for protein outliers, and that number is certain to increase.

Characterization of outliers after MMseqs2 clustering followed by selection of clusters that include 50% of the clustered proteomes provides a novel independent measure of genome assembly/annotation quality. While BUSCO can provide a more global view of proteome completeness with much less data, MMseqs2 clustering and mode-length protein outlier analysis provides a more comprehensive search for protein length errors that is not limited to a core protein set, and provides a specific validation strategy (`tfastx` searches with a mode-length protein from the cluster) to test whether a full-length protein is likely to be present. The MMseqs2 mode-length outlier strategy could be made more informative by mapping the 50% participation MMseqs2 clusters to BUSCO core proteins. Moreover, proteomes with many outliers can have very good BUSCO fragment scores (Fig. 4). Three of the nine reference *E. coli* proteomes have BUSCO fragment scores near 0, while having outlier percentages of more than 0.8%, and two of the seven reference *S. enterica* proteomes have outlier percentages greater than 0.4%, placing both sets of reference proteomes in the worst quartile of MMseqs2 clusters (Fig. 1C).

Our identification of small numbers of “missing” proteins from some proteomes, and somewhat more “missing” mode-length proteins in others, is a reminder that bacterial proteome annotation pipelines are very conservative. As a result, estimates of “core” protein sets are subject to “false-negative” errors—failures to include proteins that belong to the core orthologous set, either because the protein was truncated or it was missed altogether. Simply comparing the list of proteins annotated in a bacterial isolate does not provide the complete picture of the proteins that are present in the genome. Indeed, when the protein core of *Streptococcus agalactiae* was evaluated in the pioneering study by Tettelin et al. [4], the authors both compared the proteins that were initially annotated and also looked for “missing” proteins by searching the corresponding genome with `tfasty` (a more rigorous, but slower version of `tfastx` [16].)

Proteins annotated to be present in a proteome are almost certainly there. But proteins that appear to be “missing”, or have an unusual length, are likely to be annotation and assembly artifacts; evidence that they are in fact present in the genome, with the correct length, is often easily discovered using frameshift aware similarity searches. In this study, almost all the anomalies we see reflect technical artifacts, not biological variation.

## 5 DATA AVAILABILITY

The data for the analyses is available from Zenodo: https://doi.org/10.5281/zenodo.20208872 (alternatively FigShare: https://doi.org/10.6084/m9.figshare.32301477). The pipeline for clustering the proteomes is available from GitHub: https://github.com/g-insana/ProteomeCluster (https://doi.org/10.5281/zenodo.20208647).

The python and R scripts used to summarize the data and produce the figures is available from GitHub: https://github.com/g-insana/ett_ms (https://doi.org/10.5281/zenodo.20313420).

## 6 ACKNOWLEDGEMENTS

This work was supported by European Molecular Biology Laboratory core funds.

## SUPPLEMENTAL TABLES

**Suppl. Table 1.** MMseqs2 clustering yields.

| OSCODE | NCBI<br>taxid | proteomes<br>total (N) | clusters<br>total (N) | sequences<br>total (N) | clusters |  |  |  |  | sequences |  |  |
| --- | --- | --- | --- | --- | --- | --- | --- | --- | --- | --- | --- | --- |
|  |  |  |  |  | > 1<sup>a</sup><br>(%) | 10%<sup>b</sup><br>(N) | (%) | 50%<sup>c</sup><br>(N) | (%) | > 1<sup>a</sup><br>(%) | 10%<sup>b</sup><br>(%) | 50%<sup>c</sup><br>(%) |
| ECOLX | 562 | 22 001 | 342 137 | 106 623 057 | 47.6 | 6 716 | 2.0 | 3 927 | 1.1 | 99.8 | 88.4 | 75.2 |
| KLEPN | 573 | 17 924 | 276 163 | 95 628 119 | 42.6 | 7 121 | 2.6 | 4 686 | 1.7 | 99.8 | 93.1 | 82.7 |
| STAAU | 1280 | 13 356 | 57 134 | 35 576 073 | 50.2 | 3 562 | 6.2 | 2 383 | 4.2 | 99.9 | 95.7 | 84.8 |
| SALER | 28901 | 9 382 | 138 711 | 42 738 397 | 52.9 | 5 817 | 4.2 | 4 110 | 3.0 | 99.8 | 93.1 | 85.1 |
| STREE | 1313 | 8 865 | 34 278 | 18 195 838 | 59.3 | 2 844 | 8.3 | 1 799 | 5.2 | 99.9 | 94.3 | 82.8 |
| ACIBA | 470 | 6 712 | 97 147 | 25 023 773 | 52.6 | 4 687 | 4.8 | 3 428 | 3.5 | 99.8 | 93.3 | 85.7 |
| PSEAI | 287 | 6 367 | 149 731 | 39 731 160 | 56.1 | 7 664 | 5.1 | 5 567 | 3.7 | 99.8 | 93.8 | 86.2 |
| MYCTX | 1773 | 5 051 | 42 051 | 20 422 136 | 52.0 | 4 627 | 11.0 | 3 938 | 9.4 | 99.9 | 97.8 | 93.2 |
| LISMN | 1639 | 3 929 | 30 023 | 11 475 581 | 50.1 | 3 635 | 12.1 | 2 686 | 8.9 | 99.9 | 96.4 | 88.4 |
| ENTFC | 1352 | 2 825 | 53 444 | 7 740 206 | 49.9 | 3 859 | 7.2 | 2 298 | 4.3 | 99.6 | 92.3 | 78.6 |
| 9ENTR | 158836 | 2 540 | 90 535 | 11 852 558 | 52.9 | 6 140 | 6.8 | 3 921 | 4.3 | 99.6 | 91.9 | 81.1 |
| CLODI | 1496 | 2 152 | 37 839 | 8 016 793 | 59.9 | 4 781 | 12.6 | 3 387 | 9.0 | 99.8 | 94.7 | 87.1 |
| CAMJU | 197 | 1 845 | 20 108 | 3 091 083 | 61.1 | 2 187 | 10.9 | 1 485 | 7.4 | 99.7 | 93.5 | 84.1 |
| BURPE | 28450 | 1 632 | 51 262 | 9 753 558 | 57.6 | 7 098 | 13.8 | 5 478 | 10.7 | 99.8 | 95.4 | 89.5 |
| HELPX | 210 | 1 510 | 35 904 | 2 269 178 | 53.0 | 1 894 | 5.3 | 1 273 | 3.5 | 99.2 | 89.8 | 80.3 |
| VIBCL | 666 | 1 484 | 54 975 | 5 365 482 | 54.8 | 4 232 | 7.7 | 3 325 | 6.0 | 99.5 | 93.7 | 87.0 |
| NEIME | 487 | 1 340 | 13 393 | 2 733 748 | 64.2 | 2 858 | 21.3 | 1 800 | 13.4 | 99.8 | 95.9 | 83.4 |
| NEIGO | 485 | 1 096 | 10 889 | 2 308 163 | 56.5 | 2 720 | 25.0 | 1 927 | 17.7 | 99.8 | 97.3 | 87.4 |
| LEGNP | 446 | 937 | 16 141 | 2 869 759 | 66.7 | 4 293 | 26.6 | 2 731 | 16.9 | 99.8 | 96.5 | 85.0 |
| SHIFL | 623 | 557 | 29 904 | 2 459 171 | 50.6 | 5 141 | 17.2 | 4 065 | 13.6 | 99.4 | 95.6 | 87.5 |
| median |  |  |  |  | 53.0 | 4 460 | 8.0 | 3 356 | 5.6 | 99.8 | 94.0 | 85.0 |

<sup>a</sup> clusters with sequences from more than one proteome
<sup>b</sup> clusters containing $\geq 10\%$ of clustered proteomes
<sup>c</sup> clusters containing $\geq 50\%$ of clustered proteomes

**Suppl. Table 2.** Outlier summary.

| OSCODE | N_omes | N_clust<sup>a</sup> | N_seq<sup>a</sup> | outliers<br>(N) | outliers<br>ave. (%) | outliers<br>per cluster<br>med. (N) | outliers<br>per cluster<br>med. (%) |
| --- | --- | --- | --- | --- | --- | --- | --- |
| ECOLX | 22 001 | 3 927 | 80 189 422 | 217 722 | 0.27 | 21 | 0.10 |
| KLEPN | 17 924 | 4 686 | 79 129 907 | 172 798 | 0.22 | 20 | 0.11 |
| STAAU | 13 356 | 2 383 | 30 171 995 | 81 598 | 0.27 | 12 | 0.09 |
| SALER | 9 382 | 4 110 | 36 361 199 | 102 350 | 0.28 | 12 | 0.14 |
| STREE | 8 865 | 1 799 | 15 064 785 | 104 617 | 0.69 | 9 | 0.10 |
| ACIBA | 6 712 | 3 428 | 21 452 377 | 68 947 | 0.32 | 9 | 0.15 |
| PSEAI | 6 367 | 5 567 | 34 248 121 | 104 924 | 0.31 | 6 | 0.09 |
| MYCTX | 5 051 | 3 938 | 19 036 113 | 95 834 | 0.50 | 6 | 0.12 |
| LISMN | 3 929 | 2 686 | 10 140 993 | 8 017 | 0.08 | 1 | 0.03 |
| ENTFC | 2 825 | 2 298 | 6 082 249 | 23 376 | 0.38 | 2 | 0.07 |
| 9ENTR | 2 540 | 3 921 | 9 608 252 | 12 779 | 0.13 | 1 | 0.04 |
| CLODI | 2 152 | 3 387 | 6 978 669 | 34 294 | 0.49 | 3 | 0.14 |
| CAMJU | 1 845 | 1 485 | 2 599 303 | 6 085 | 0.23 | 2 | 0.11 |
| BURPE | 1 632 | 5 478 | 8 726 277 | 34 731 | 0.40 | 0 | 0.00 |
| HELPX | 1 510 | 1 273 | 1 822 489 | 6 145 | 0.34 | 3 | 0.20 |
| VIBCL | 1 484 | 3 325 | 4 669 404 | 20 042 | 0.43 | 2 | 0.14 |
| NEIME | 1 340 | 1 800 | 2 280 328 | 10 556 | 0.46 | 1 | 0.07 |
| NEIGO | 1 096 | 1 927 | 2 016 990 | 10 029 | 0.50 | 0 | 0.00 |
| LEGNP | 937 | 2 731 | 2 439 719 | 3 700 | 0.15 | 0 | 0.00 |
| SHIFL | 557 | 4 065 | 2 151 969 | 16 090 | 0.75 | 1 | 0.18 |
| median | 2 682 | 3 356 | 9 167 264 | 28 835 | 0.30 | 2 | 0.10 |

<sup>a</sup> in 50% participation clusters

## SUPPLEMENTAL FIGURES

**Suppl. Fig. 1.**
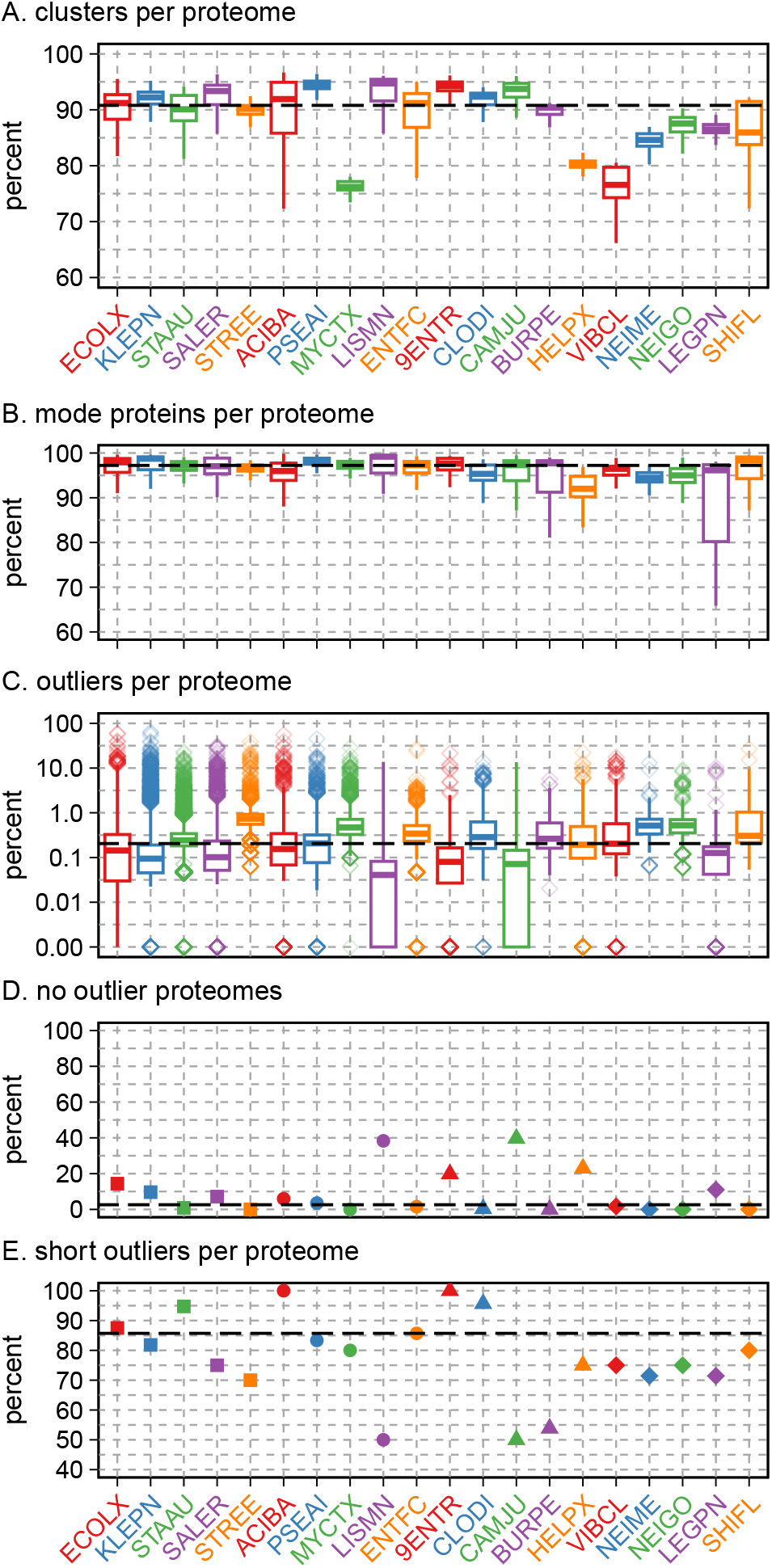
Proteome cluster content, quality, and error types, across the 20 bacterial species. This figure displays the same measures of quality as Fig. 1, with each bacteria showing the properties of proteomes, rather than clusters. (A) Clusters from each proteome divided by the median number of proteins in each of the 20 bacterial species. (B) The percent of the proteins in each proteome that match the mode length of the cluster. (C) The percent of outliers from each bacterial proteome. (D) The percent of bacterial proteomes with no outliers. (E) Percent of outliers per proteome that are short. Dashed lines show the median of the medians for the 20 bacteria.

**Suppl. Fig. 2.**
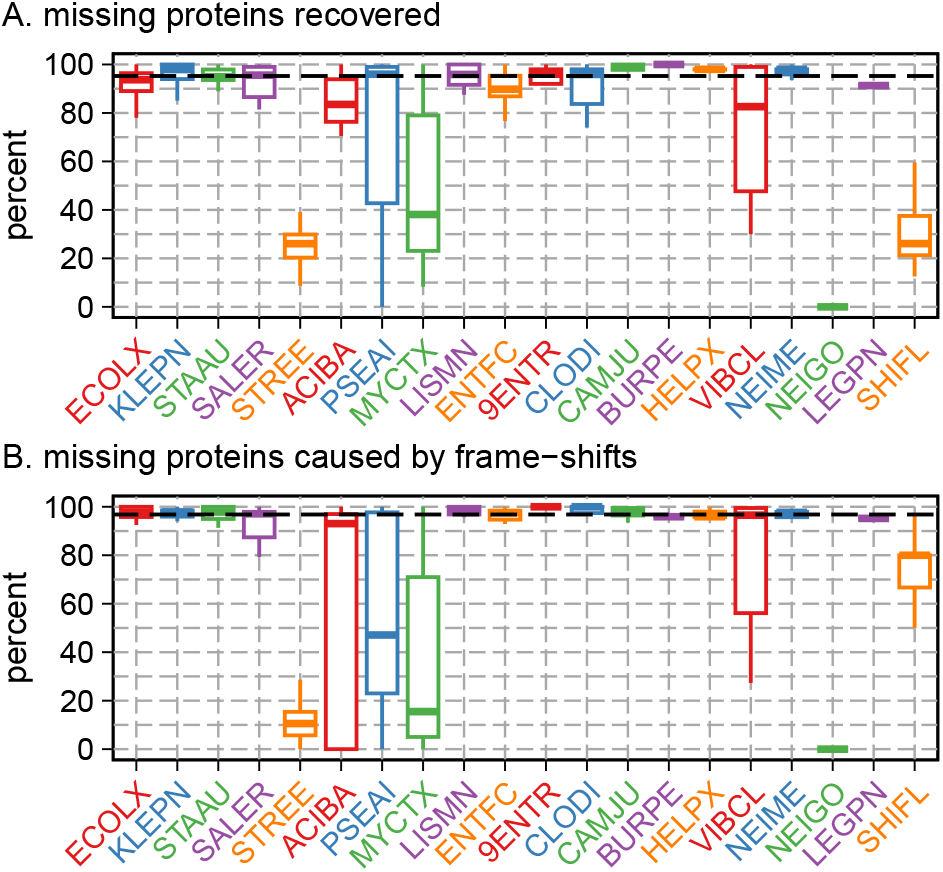
Recovery of mode-proteins in genomes annotated to not participate in a cluster. Up to 50 randomly selected mode-length proteins from missing clusters were compared to up to 20 proteomes that did contribute to the respective clusters. Results for 20 bacteria are shown, but NEIGO was not analyzed because no NEIGO proteomes contributed to fewer than 2/3 the clusters for that bacteria. (A) The percent of full-length *>*90% identity alignments that were found for each non-contributing genome. The dashed line indicates the median recovery (95.2%) across the 20 proteomes. (B) The percent of alignments that included a frame-shift (median 96.8%, dashed line).

**Suppl. Fig. 3.**
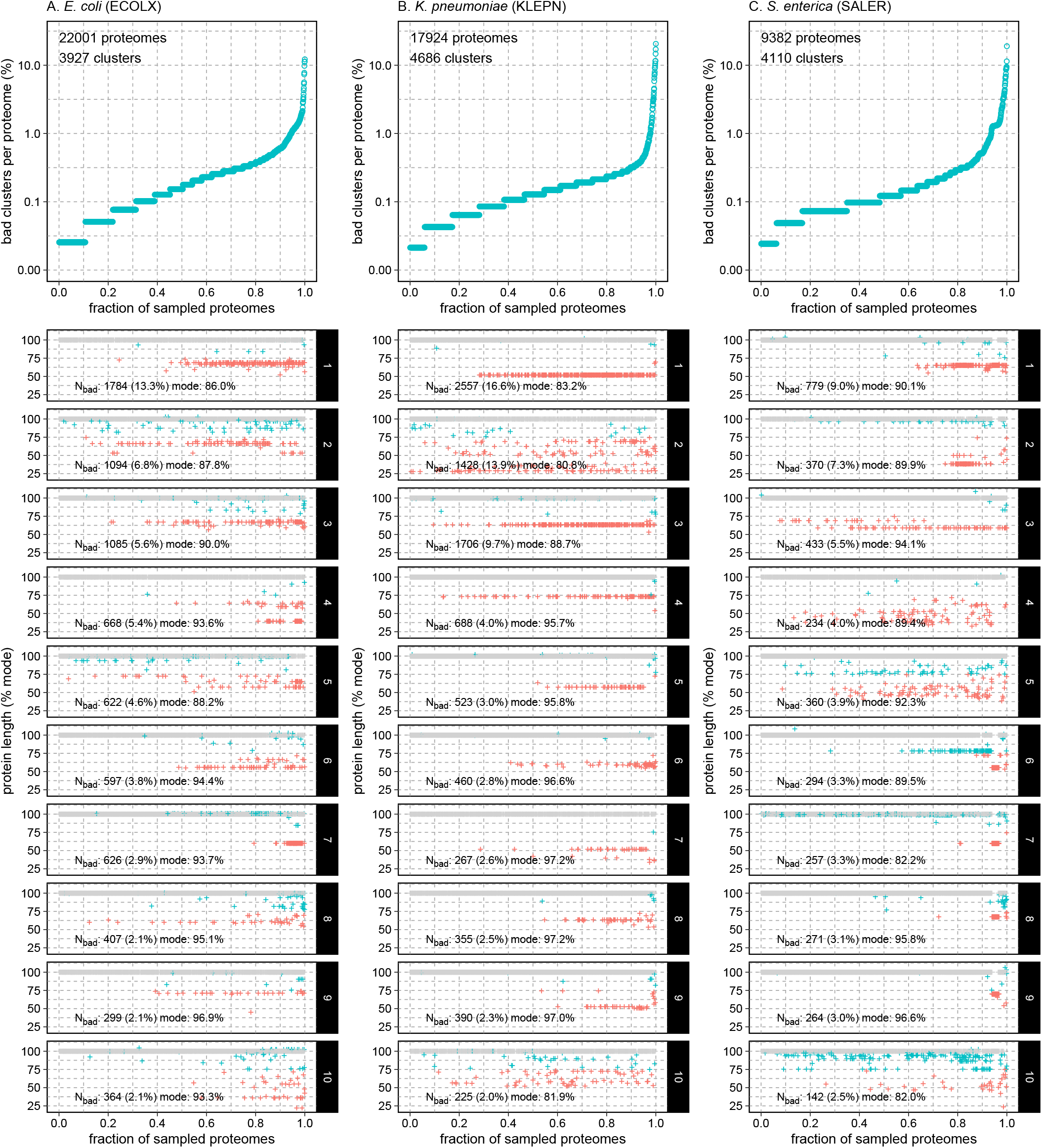
Protein lengths across sampled proteomes. Clusters from three bacteria: (A) *E. coli*, (B) *K. pneumoniae*, and (C) *S. enterica* are shown. Each top panel (A, B, C) shows the total number of bad clusters per proteome for a sampled set of up to 1000 proteomes. The bottom 10 panels show the distribution of protein lengths across the sampled proteomes for 10 randomly selected clusters. N_*bad*_ shows the number of outlier proteins across all the proteomes in the cluster, and the percent (%) of outlier proteins in the cluster. “mode” reports the fraction of proteins in the cluster with the mode length. In the bottom 10 panels, the + symbols show the normalized length (% of the mode length) for every protein in the cluster that was seen in the sampled proteomes. Grey + symbols have the mode length, cyan + are *>*0.75 mode-length or *<* 1.33. Magenta + symbols denote outliers *<*0.75 mode-length.

**Suppl. Fig. 4.**
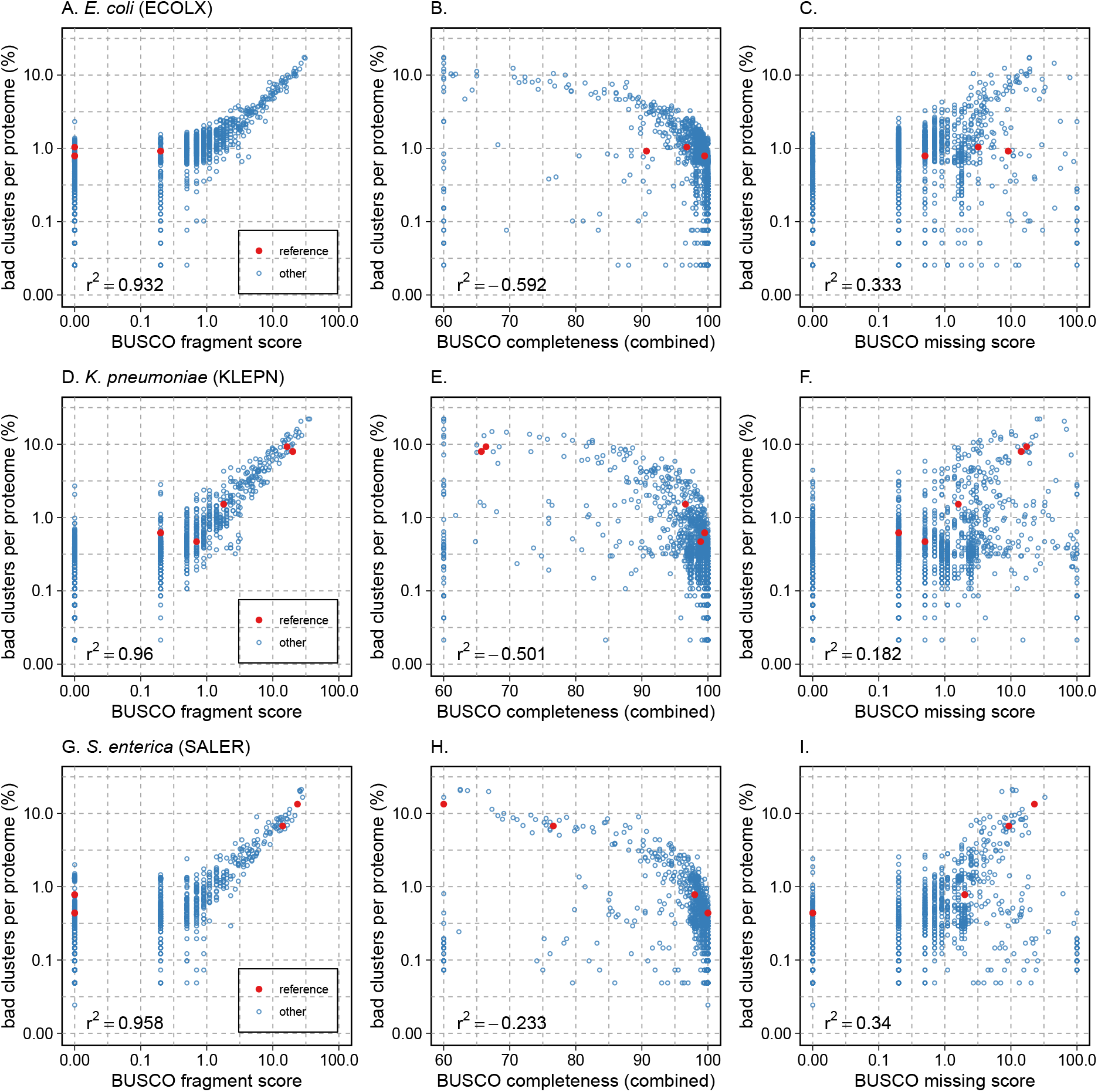
Comparison of outlier-frequency (%) to BUSCO measures of proteome quality for three of the proteomes shown in Fig. 3, *E. coli, K. pneumoniae*, and *S. enterica*. Correlation coefficients (*r*^2^) were calculated for the untransformed (linear) relations between BUSCO measure and the percentage of bad clusters in each individual proteome isolate. The BUSCO completeness (combined) score is plotted; the completeness (single) score is indistinguishable from the completeness (combined) score.

**Suppl. Fig. 5.**
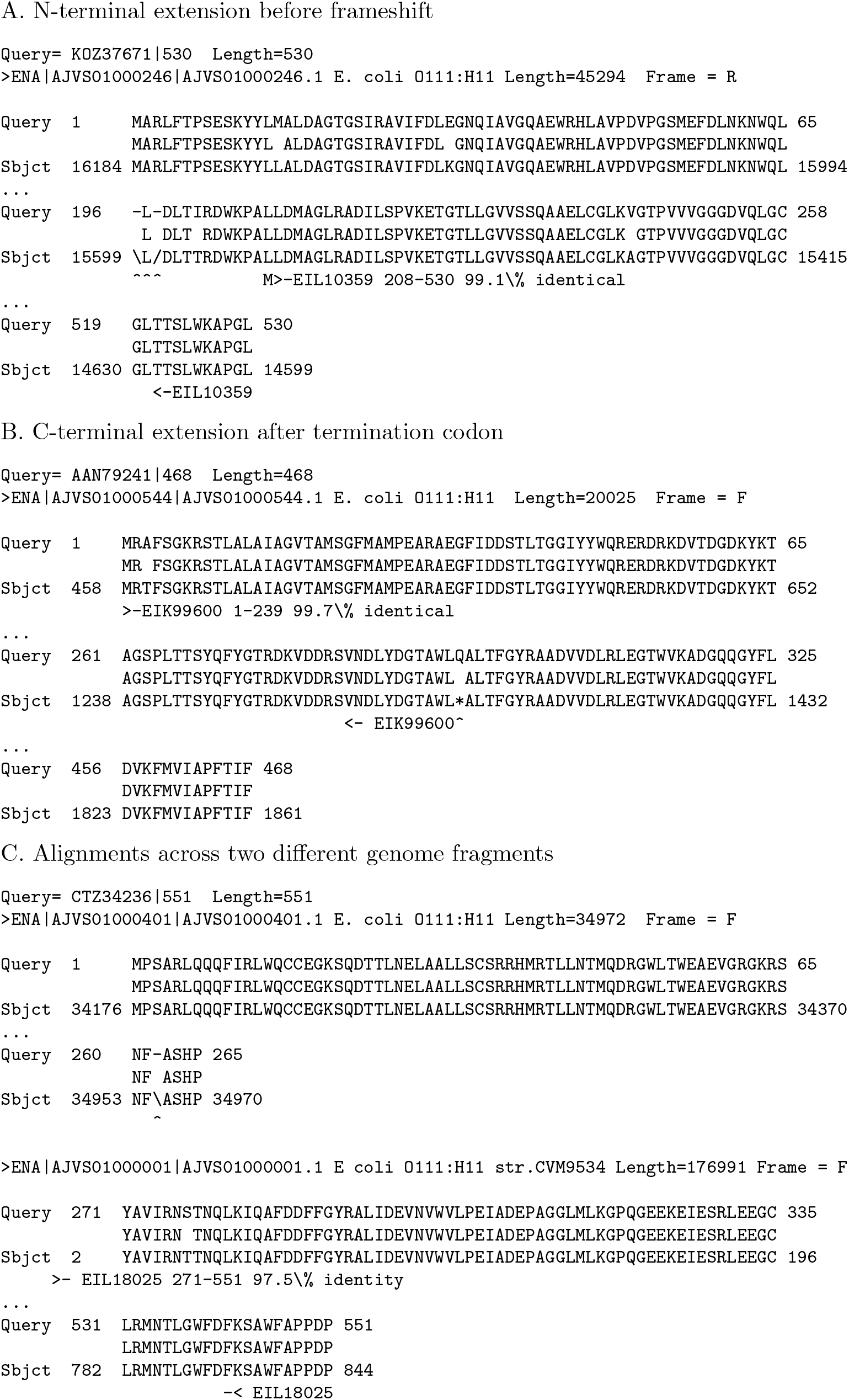
Alignment of mode length (full-length) proteins to a genome that produced short proteins.

**Suppl. Fig. 5.**
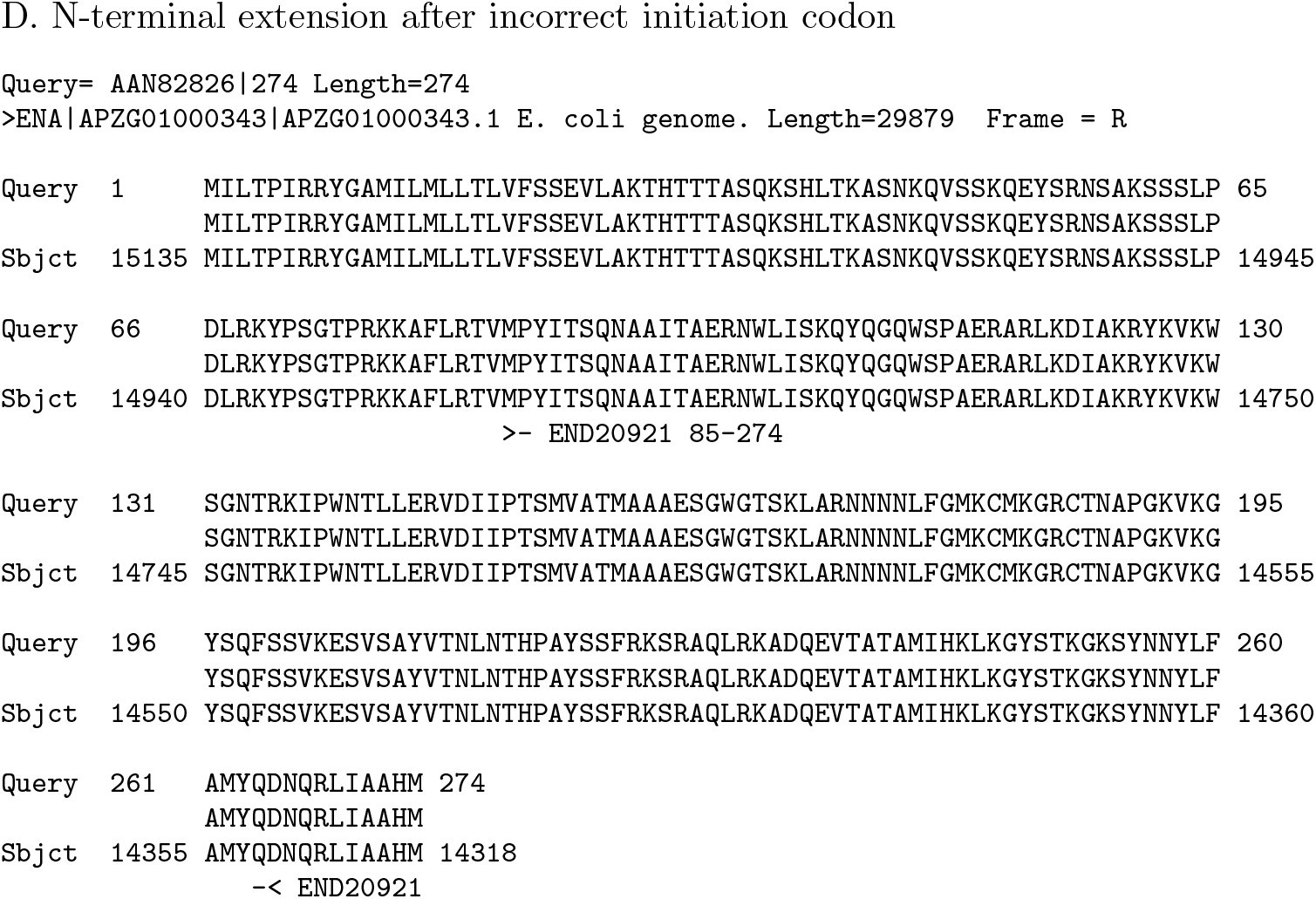
Alignment of mode length (full-length) proteins to a genome that produced short proteins. Only the boundaries of the alignment between the long (mode-length) protein, and the boundaries of the alignment to the shorter protein annoted from the proteom are shown. (A) Alignment of mode length protein `KOZ37671` (530 amino-acids) to `GCA 000263935.1`, which is annotated to produce `EIL10359` (323 aa). (B) Alignment of `AAN79241` (551 aa) to `GCA 000263935.1`, which is annotated to produce `EIK99600` (239 aa). (C) Alignment of `CTZ34236` (551 aa) to `GCA 000263935.1`, annotated to produce `EIL18025` (281 aa). (D) Alignment of `AAN82826` (274 aa) to `GCA 000357805.1`, annotated to produce `END20921` (190 aa). The ‘*\*’ and’`/` ‘characters indicate frameshifts, the’ * ‘ character indicates a termination codon.

**Suppl. Fig. 6.**
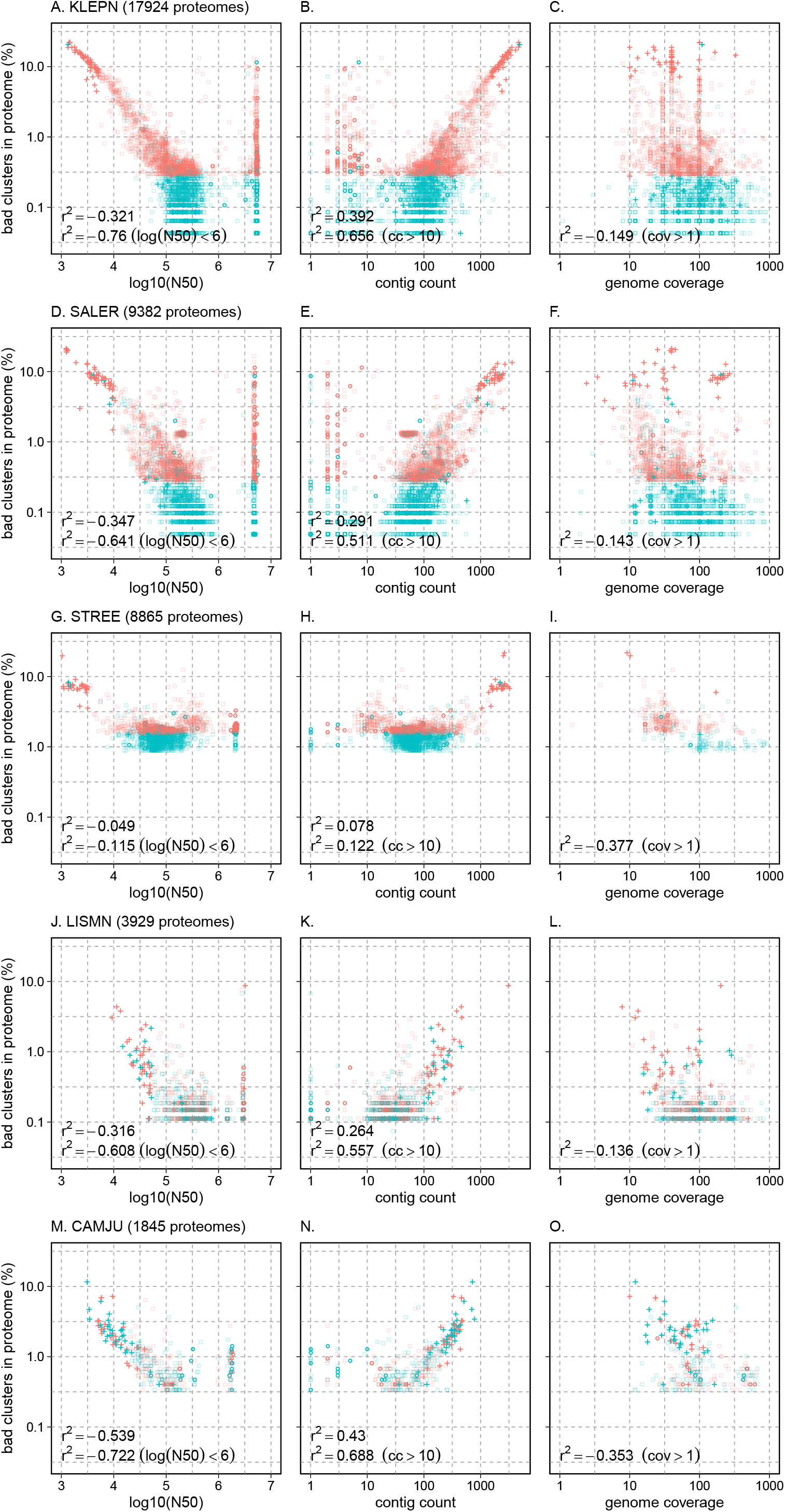
Comparison of outlier-frequency (%) to median contig length (log10(N50)), contig count, and sequence coverage for *K. pneumoniae* (KLEPN), *S. enterica* (SALER), *S. pneumoniae* (STREE) and *L. monocytogenes* (LISMN) and *C. jejuni* (CAMJU). Symbols are plotted as in Fig. 6. Red symbols plot the worst proteomes by percentage bad clusters; blue symbols plot a sample of all proteomes.

**Suppl. Fig. 7.**
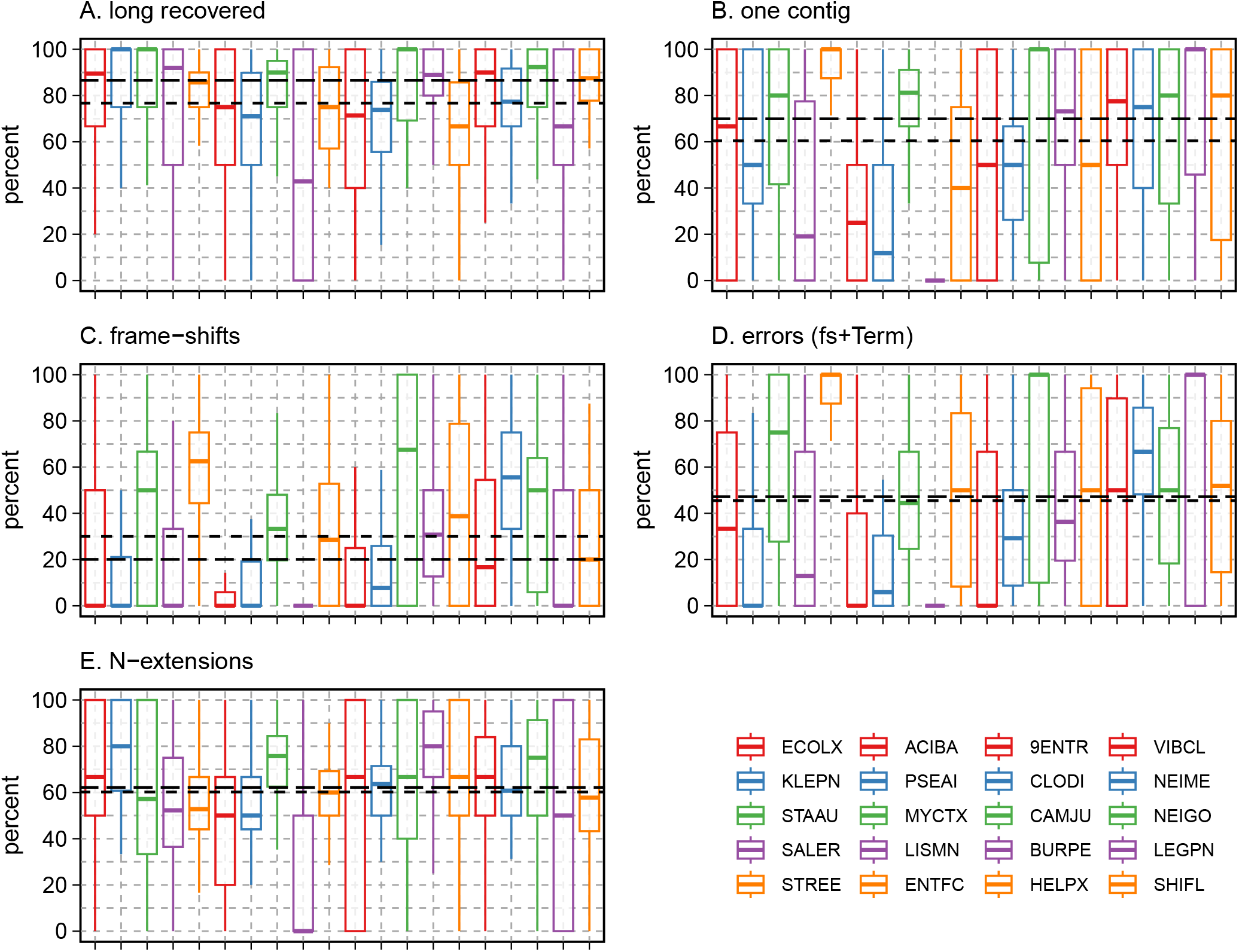
Recovery of full-length (mode-length) proteins from “short” proteomes. Mode-length proteins were aligned with up to 100 randomly sampled proteomes with at least 5 short-outlier proteins from each of the 20 bacteria surveyed to identify full-length (“mode-length”) alignments. (A) The percent of alignments that recovered a full-length mode-length alignment in the bacterial genome that produced the short protein. (B) The percent of recovered alignments found in a single contig. (C) The percent of recovered alignments that included a frame-shift. (D) The percent of recovered alignments that included a frame-shift or termination codon. (E) The percent of recovered alignment extensions towards the N-terminal of the longer protein. In each panel, the shorter-dashed line indicates the median of the average percentage across the 20 bacteria, while the longer-dashed line indicates the median of the medians for the 20 bacteria.

